# Sex-specific stress-related behavioral phenotypes and central amygdala dysfunction in a mouse model of 16p11.2 microdeletion

**DOI:** 10.1101/2020.11.12.380501

**Authors:** Jacqueline Giovanniello, Sandra Ahrens, Kai Yu, Bo Li

## Abstract

Substantial evidence indicates that a microdeletion on human chromosome 16p11.2 is linked to neurodevelopmental disorders including autism spectrum disorders (ASD). Carriers of this deletion show divergent symptoms besides the core features of ASD, such as anxiety and emotional symptoms. The neural mechanisms underlying these symptoms are poorly understood. Here we report mice heterozygous for a deletion allele of the genomic region corresponding to the human 16p11.2 microdeletion locus (i.e., the ‘*16p11.2 del*/+ mice’) have sex-specific anxiety-related behavioral and neural circuit changes. We found that female, but not male *16p11.2 del*/+ mice showed enhanced fear generalization – a hallmark of anxiety disorders – after auditory fear conditioning, and displayed increased anxiety-like behaviors after physical restraint stress. Notably, such sex-specific behavioral changes were paralleled by an increase in activity in central amygdala neurons projecting to the globus pallidus in female, but not male *16p11.2 del*/+ mice. Together, these results reveal female-specific anxiety phenotypes related to 16p11.2 microdeletion syndrome and a potential underlying neural circuit mechanism. Our study therefore identifies previously underappreciated sex-specific behavioral and neural changes in a genetic model of 16p11.2 microdeletion syndrome, and highlights the importance of investigating female-specific aspects of this syndrome for targeted treatment strategies.

## Introduction

As of 2018, the Autism and Developmental Disabilities Monitoring (ADDM) Network of the Center for Disease Control (CDC) estimated that approximately one in 59 children age eight and younger are currently diagnosed with autism spectrum disorders (ASD) (Baio et al., 2018). ASD is a spectrum of neurodevelopmental conditions defined by two major diagnostic criteria: “persistent deficits in social communication and social interaction across multiple contexts” and “restricted, repetitive patterns of behavior, interests, or activities” (*Diagnostic and Statistical Manual of Mental Disorders, DSM-5*, 2013). Diagnoses of ASD often include supplemental association with intellectual disability, catatonia, other defined neurodevelopmental, mental, behavioral disorders, and/or a known medical, genetic, or environmental factor. Furthermore, patients with ASD are commonly diagnosed with one or more comorbid conditions including intellectual disability (Howlin, 2000; Schwartz & Neri, 2012; Tonnsen et al., 2016), attention deficit-hyperactivity disorder (Antshel et al., 2014, 2016; Antshel & Russo, 2019; Jang et al., 2013), obsessive compulsive disorder (Leyfer et al., 2006; Postorino et al., 2017), anxiety (Brookman-Frazee et al., 2018; Gotham et al., 2013; White et al., 2009), and depression (Andersen et al., 2015; Davidsson et al., 2017; Gotham et al., 2013; Matson & Cervantes, 2014), and are at increased risk for suicidality, particularly among females (T Hirvikoski et al., 2019; Tatja Hirvikoski et al., 2016; Kirby et al., 2019).

Despite the heterogeneity in ASD features, one major consistency is its sex bias in diagnoses. It is well documented that ASD is about 4 times more common in males than in females with an exception for x-linked syndromes, such as Rett Syndrome which is more common in females (Fombonne, 2002). There is significant evidence of divergence among core symptoms of ASD based on sex. Specifically, many studies have found reduced severity of repetitive and or stereotyped behaviors in females than in males (Baron-Cohen, 2009; Beggiato et al., 2017; Knickmeyer et al., 2008; Kopp et al., 2010; Szatmari et al., 2012). In contrast, females show different social impairments compared with males (Beggiato et al., 2017; Dean et al., 2017; Head et al., 2014; Hiller et al., 2014; Werling & Geschwind, 2013). These tend toward more internalizing symptoms and emotional disturbance (Horiuchi et al., 2014; Kreiser & White, 2014; Rynkiewicz et al., 2016; Rynkiewicz & Łucka, 2018; Solomon et al., 2012). Females with ASD also show increased risk of eating disorders (Kalyva, 2009), sensory impairments (Lai et al., 2014), sleep disturbances (Hartley & Sikora, 2009), epilepsy and learning disorders (Giarelli et al., 2010). It has been suggested that females may “camouflage” their autism phenotypes better than males owing to fewer social impairments and better executive functioning (Bölte et al., 2011), as well as reduced externalizing symptoms (Werling & Geschwind, 2013). One way that emotional phenotypes often manifest, is as anxiety disorders. In the general population, females have an increased prevalence of stress-related disorders such as anxiety, depression, and PTSD (Breslau, 2002; Kessler et al., 1995; Olff, 2017; Tolin & Foa, 2006). Therefore, it is possible that anxiety-like phenotypes may present differently in males and females with ASD.

A major limitation of much of the research in ASD has been its emphasis on males. This is not exclusive to ASD research as most research is done in males (Hughes, 2007). Among neuroscience studies in general, the sex bias of human subjects is approximately 5.5 males for every female and with a ratio much higher among animal studies (Beery & Zucker, 2011). This bias precludes our understanding of autism in females and limits our development of effective treatment strategies. Therefore, we sought to examine whether sex differences exist in stress-related behaviors in a mouse model of ASD. To this end, we utilized a model that mimics a microdeletion on human chromosome 16p11.2. Notably, this deletion is one of the most common genetic variations found in ASD, accounting for ~1% of ASD cases (Chen et al., 2017; Kumar et al., 2007; Marshall et al., 2008; Sanders et al., 2011; Sebat et al., 2007; Weiss et al., 2008). Patients with this deletion show repetitive behaviors, hyperactivity, intellectual disability, motor and speech/language delay, and anxiety symptoms (Al-Kateb et al., 2014; Bijlsma et al., 2009; Fernandez et al., 2010; Shinawi et al., 2010; Steinman et al., 2016). Of note, individuals carrying the 16p11.2 deletion, including those non-ASD carriers, are often diagnosed as having anxiety disorders or other mood disorders (Zufferey et al., 2012).

The mouse model we used was generated by Horev et al. (Horev et al., 2011), and is one of three independently generated mouse genetic models that mimic the 16p11.2 microdeletion (Arbogast et al., 2016; Horev et al., 2011; Portmann et al., 2014). These models, which were created by deleting largely similar genomic intervals in mouse chromosome 7 corresponding to the syntenic 16p11.2 microdeletion region in humans, exhibit overlapping phenotypes (Arbogast et al., 2016; Horev et al., 2011; Portmann et al., 2014). In particular, heterozygous deletion mice – hereafter referred to as *16p11.2 del/+* mice – in each of these lines share basic phenotypes such as low body weight and perinatal mortality, and, importantly, also show behavioral phenotypes related to the symptoms of human 16p11.2 microdeletion carriers. These phenotypes include increased locomotor activity, stereotyped and repetitive behaviors, sleep deficits, recognition memory deficits, reward learning deficits, and social deficits (Angelakos et al., 2017; Arbogast et al., 2016; Grissom et al., 2017; Horev et al., 2011; Portmann et al., 2014; Rein & Yan, 2020; Walsh et al., 2018; Yang, Lewis, et al., 2015; Yang, Mahrt, et al., 2015).

A few studies examined the *16p11.2 del/+* mice for anxiety or fear-related behaviors, but with mixed results. When tested in the open field (OPF) test and elevated plus maze (EPM) test, assays conventionally used to assess ‘anxiety’ in rodents, these mice appear not different from wildtype (WT) mice (Arbogast et al., 2016; Lynch et al., 2020; Yang, Lewis, et al., 2015), (however, see (Pucilowska et al., 2015)). The *16p11.2 del/+* mice were also examined in fear conditioning paradigms. One study shows that the *16p11.2 del/+* mice have impaired contextual fear conditioning (Tian et al., 2015), whereas other studies show that the *16p11.2 del/+* mice have normal contextual fear conditioning and normal visually cued fear conditioning (Lynch et al., 2020; Yang, Lewis, et al., 2015).

Recent studies indicate that environmental factors can exacerbate ASD symptomatology and impairments in cognitive and adaptive behaviors in 16p11.2 deletion carriers (Hudac et al., 2020), and *16p11.2 del/+* mice show altered coping in response to stress compared with wildtype littermates (Panzini et al., 2017; Yang, Lewis, et al., 2015). In light of these findings and studies showing males and females can exhibit very different behavioral responses to threats or stress (Dalla & Shors, 2009; Gruene et al., 2015), we reasoned that under a stressful situation *16p11.2 del/+* mice may exhibit sex-specific behavioral changes. However, a potential sex-specific effect of the *16p11.2* deletion on anxiety or fear-related behaviors in mice has not been considered until recently (Lynch et al., 2020). Furthermore, only simple assays, such as OPF and EPM tests, have been used to assess “baseline anxiety” in *16p11.2 del/+* mice, which may not be sufficient to reveal potential changes in anxiety or fear processing in response to stress in these mice.

To address these issues, in the current study we examined anxiety-related behaviors under different stress conditions in both male and female *16p11.2 del/+* mice and their wild type littermates. We found that female, but not male *16p11.2 del/+* mice showed enhanced fear generalization, a hallmark of anxiety disorders (Dunsmoor & Paz, 2015), after auditory fear conditioning. Furthermore, although at baseline *16p11.2 del/+* mice were not different from their wildtype littermates in the EPM test, consistent with previous studies (Arbogast et al., 2016; Lynch et al., 2020; Yang, Lewis, et al., 2015), we found that female, but not male *16p11.2 del/+* mice showed enhanced anxiety in the EPM after acute restraint stress. Lastly, we found that these sex-specific behavioral changes were paralleled by an increase in activity in the central amygdala – a major limbic structure that regulates anxiety in rodents and primates (Ahrens et al., 2018; Fox et al., 2012; Shackman & Fox, 2016) – of female, but not male *16p11.2 del/+* mice. Together, our work suggests that 16p11.2 microdeletion differentially affects males and females and may disproportionally disrupt stress-regulation brain functions in females. These findings provide insight into understanding how ASD may present differently in females at behavioral and neuronal levels, and raise the question of whether changes to treatment and diagnostic strategies based on sex should be considered.

## Methods

### Animals

To breed *16p11.2 del/+* mice, we used *16p11.2 del/+* male mice (Stock Number: 013128) and C57/B6 female mice purchased from the Jackson Laboratory, or similar breeders obtained from Pavel Osten’s lab at Cold Spring Harbor Laboratory (CSHL). Breeders were housed with a cardboard bio-hut under a 12-hour light/dark cycle (7 am to 7 pm light) with food and water available *ad libitum*. As *16p11.2 del/+* mice exhibit postnatal lethality (Horev et al., 2011), in breeding cages only, standard rodent chow (LabDiet) was supplemented with DietGel^®^ Boost (ClearH2O), a high calorie liquid diet that increased survival of *16p11.2 del*/+ pups. Pups were weaned at 3 weeks of age and group housed with wildtype littermates. Mice were genotyped for 16p11.2 microdeletion between 4-8 weeks of age with primers provided by Alea Mills’ lab at CSHL.

Mice of 2-4 months old were used for all behavioral experiments. Mice of 6-10 weeks old were used for all electrophysiology experiments. All experimental mice were housed under a 12-h light/dark cycle (7 a.m. to 7 p.m. light) in groups of 2-5 animals, containing both *16p11.2 del/+* mice and their wildtype littermates. Food and water were available *ad libitum*. All behavioral experiments were performed during the light cycle. Littermates were randomly assigned to different groups prior to experiments. All experimental procedures were approved by the Institutional Animal Care and Use Committee of CSHL and performed in accordance to the US National Institutes of Health guidelines.

### Behavioral tasks

#### Auditory fear conditioning

We followed standard procedures for classical auditory fear conditioning (Li et al., 2013; Penzo et al., 2014, 2015; Yu et al., 2017). Briefly, mice were initially handled and habituated to a conditioning cage, which was a Mouse Test Cage (18 cm x 18 cm x 30 cm) with an electrifiable floor connected to a H13-15 shock generator (Coulbourn Instruments, Whitehall, PA). The Test Cage was placed inside a sound attenuated cabinet (H10-24A; Coulbourn Instruments). Before each habituation and conditioning session, the Test Cage was wiped with 70% ethanol. The cabinet was illuminated with white light during habituation and conditioning sessions.

During habituation, two 4-kHz 75-dB tones and two 12-kHz 75-dB tones, each of which was 30s in duration, were delivered at variable intervals within an 8-minute session. During conditioning, mice received three presentations of the 4-kHz tone (conditioned stimulus; CS+), each of which co-terminated with a 2-s 0.7-mA foot shock (unless otherwise stated), and three presentations of the 12-kHz tone (CS–), which were not paired with foot shocks. The CS+ and CS– were interleaved pseudo-randomly, with variable intervals between 30 and 90 s within a 10-minute session. The test for fear memory (retrieval) was performed 24 h following conditioning in a novel context, where mice were exposed to two presentations of CS+ and two presentations of CS– (>120 s inter-CS interval). The novel context was a cage with a different shape (22 cm x 22 cm x 21 cm) and floor texture compared with the conditioning cage, and was illuminated with infrared light. Prior to each use the floor and walls of the cage were wiped clean with 0.5% acetic acid to make the scent distinct from that of the conditioning cage.

Animal behavior was videotaped with a monochrome CCD-camera (Panasonic WV-BP334) at 3.7 Hz and stored on a personal computer. The FreezeFrame software (Coulbourn Instruments) was used to control the delivery of both tones and foot shocks. Freezing behavior was analyzed with FreezeFrame software (Coulbourn Instruments). Baseline freezing levels were calculated as the average freezing during the first 100 s of the session before any stimuli were presented, and freezing to the auditory stimuli was calculated as the average freezing during the tone presentation. The average of the freezing responses to two CS+ or CS– presentations during retrieval was used as an index of fear. Discrimination Index was calculated as the difference between freezing to the CS+ and CS–, normalized by the sum of freezing to both tones.

#### Shock sensitivity test

Animals were placed in a conditioning Test Cage in a lit, sound attenuated box, as in the fear conditioning experiments, and received two presentations each of 0.2, 0.4, 0.6, 0.8, and 1.0 mA shocks with an inter-shock interval of 30 seconds. Animals were monitored with a monochrome CCD camera (Panasonic WV-BP334) at 4 Hz, and tracked and analyzed using Ethovision XT 5.1 (Noldus Information Technologies) to extract distance traveled and movement velocity during the 2s time window of each shock presentation.

#### Acute physical restraint stress

For stress susceptibility experiments, animals underwent a standard protocol of acute physical restraint as described previously (K. Kim & Han, 2006). Mice were immobilized in a well-ventilated 50 mL conical tube for two hours in a dark, sound attenuated chamber. Males and females were kept in separate chambers. Animals were then tested on the EPM 24 hours after the end of the restraint session.

#### Elevated plus maze test

The elevated plus maze (EPM) test apparatus was constructed from white Plexiglas and consisted of two open arms without walls (30 cm long and 5 cm wide) and two arms enclosed by 15.25 cm high non-transparent walls. The arms were extended from a central platform (5 cm x 5 cm), and were arranged such that the identical arms were opposite to each other. The maze was raised to a height of 50 cm above the floor with an overhead light. At the start of the session, animals were placed in the center zone and allowed to explore the maze for 10 minutes in the absence of the experimenter, while their behavior was videotaped using a monochrome CCD camera (Panasonic WV-BP334) at 4 Hz. The resulting data was analyzed using Ethovision XT 5.1 (Noldus Information Technologies). Parameters assessed were total distance travelled, velocity, time spent in the open arms, number of entries to the open arms, and latency to the first entry into an open arm. The maze was thoroughly cleaned with 70% ethanol in between subjects.

#### Auditory discrimination test

Mice were first trained in an auditory two-alternative choice (2-AC) procedure as previously described (Ahrens et al., 2015). Briefly, mice initiated each trial by poking their nose into the center port of a three-port operant chamber. After a silent delay of random duration (200–300 ms, uniformly distributed), a frequency-modulated target sound was presented. The carrier frequency of the target indicated to the animal which of the two side ports would provide 10 μl of water reward. For a target carrier frequency of 4kHz, reward was available only at the left port. For a target of 12 kHz, reward was provided at the right port. Mice were only rewarded in trials in which they chose the correct port as their first choice. Sound intensity was set at 60 dB-SPL, and sound duration was 100 ms. The modulation frequency was set to 15 Hz. Incorrect choices were punished by a 4s timeout and a white noise presentation.

After mice reached a performance level of 70% in the 2-AC task, they were tested for auditory discrimination. Mice initiated a trial by a nose poke into the center port. After a silent delay of random duration (200-300 ms), a frequency modulated sound was presented for 100 ms. The frequency of the sound was randomly selected from a group of eight frequencies (4, 4.68, 5.48, 6.4, 7.49, 8.77, 10.26 and 12 kHz). These frequencies were chosen such that they were equidistant from each other on the logarithmic (Log2) scale. All frequencies less than 6.9 kHz (the geometric mean of 4 and 12 kHz) were rewarded if the mouse chose the left water port, and those greater than 6.9 kHz were rewarded with water in the right water port. The volume of the water reward was 5 μl to ensure that the mice performed sufficient number of trials for each of the frequencies. Data from five consecutive sessions were collected (250-350 trials per session). Responses of each mouse to the eight sound frequencies was transformed into the percentage of ‘proportion right choice’, which is the percentage of the trials in which the mouse chose the water port on the right side. These data were fitted using the following logistic function(Ahrens et al., 2015; Gilchrist et al., 2005):

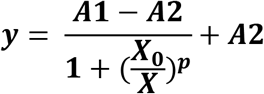

where *X*_0_ represents the median threshold and *p* determines the slope of the curve; *A1* and *A2* are the upper and lower bounds of the equation, respectively. A sigmoidal psychometric curve was thus generated. The median threshold *X*_0_ and parameter *p* of this curve were then obtained for each animal, and the data were pooled for each group.

### Stereotaxic Surgery

Standard surgical procedures were followed for stereotaxic injection (Li et al., 2013; Penzo et al., 2015; Yu et al., 2016, 2017). Briefly, mice were anesthetized with isoflurane (3% at the beginning and 1% for the rest of the surgical procedure), and positioned in a stereotaxic injection frame (myNeuroLab.com). A digital mouse brain atlas was linked to the injection frame to guide the identification and targeting (Angle Two Stereotaxic System, myNeuroLab.com). The injection was performed at the following stereotaxic coordinates for GPe: −0.46 mm from Bregma, 1.85 mm lateral from the midline, and 3.79 mm vertical from skull surface.

For injection of the retrograde tracer, we made a small cranial window (1–2 mm^2^), through which the tracer (~0.3 μl) was delivered via a glass micropipette (tip diameter, ~5 μm) by pressure application (5–20 psi, 5–20 ms at 0.5 Hz) controlled by a Picrospritzer III (General Valve) and a pulse generator (Agilent). During the surgical procedure, mice were kept on a heating pad maintained at 35°C and were brought back to their home-cage for post-surgery recovery and monitoring. Subcutaneous Metacam (1-2 mg kg–1 meloxicam; Boehringer Ingelheim Vetmedica, Inc.) was given post-operatively for analgesia and anti-inflammatory purposes.

The retrograde tracer cholera toxin subunit B (CTB) conjugated with Alexa Fluor™ 555 (CTB-555) was purchased from Invitrogen, Thermo Fisher Scientific (Waltham, Massachusetts, USA). CTB was used at a concentration of 1mg/ml in phosphate-buffered saline and kept at −20°C.

### *In vitro* electrophysiology

For the *in vitro* electrophysiology experiments, mice were anaesthetized with isoflurane and perfused intracardially with 20 mL ice-cold artificial cerebrospinal fluid (ACSF) (118 mM NaCl, 2.5 mM KCl, 26.2 mM NaHCO_3_, 1 mM NaH_2_PO_4_, 20 mM glucose, 2 mM MgCl_2_ and 2 mM CaCl_2_, pH 7.4, gassed with 95% O_2_ and 5% CO_2_). Mice were then decapitated and their brains quickly removed and submerged in ice-cold dissection buffer (110.0 mM choline chloride, 25.0 mM NaHCO_3_, 1.25 mM NaH_2_PO_4_, 2.5 mM KCl, 0.5 mM CaCl_2_, 7.0 mM MgCl_2_, 25.0 mM glucose, 11.6 mM ascorbic acid and 3.1mM pyruvic acid, gassed with 95% O2 and 5% CO2). Coronal sections of 300 μm containing the central amygdala (CeA) were cut in dissection buffer using a HM650 Vibrating-blade Microtome (Thermo Fisher Scientific). Slices were immediately transferred to a storage chamber containing ACSF at 34 °C. After 40 min recovery time, slices were transferred to room temperature (20–24°C) and perfused with gassed ACSF constantly throughout recording.

Whole-cell patch clamp recording was performed as previously described (Li et al., 2013). Briefly, recording from CeA neurons was obtained with Multiclamp 700B amplifiers and pCLAMP 10 software (Molecular Devices, Sunnyvale, California, USA), and was visually guided using an Olympus BX51 microscope equipped with both transmitted and epifluorescence light sources (Olympus Corporation, Shinjuku, Tokyo, Japan). The external solution was ACSF. The internal solution contained 115 mM cesium methanesulfonate, 20 mM CsCl, 10 mM HEPES, 2.5 mM MgCl_2_, 4 mM Na2ATP, 0.4 mM Na_3_GTP, 10 mM sodium phosphocreatine and 0.6 mM EGTA (pH 7.2). Miniature excitatory post-synaptic currents (mEPSCs) were recorded at −70 mV with bath application of 100 μM GABA antagonist, picrotoxin (PTX), and 1 μM sodium channel blocker, tetrodotoxin (TTX). The internal solution contained 115 mM cesium methanesulphonate, 20 mM CsCl, 10 mM HEPES, 2.5 mM MgCl_2_, 4 mM Na_2_-ATP, 0.4 mM Na_3_GTP, 10 mM Na-phosphocreatine and 0.6 mM EGTA (pH 7.2, 290 mOsm). Data was collected in gap-free mode in pClamp 10 (Molecular Devices) for 5 minutes at room temperature and analyzed using Mini Analysis Program (Synaptosoft). For recordings on CeA neurons projecting to the GPe, CTB-555 was injected into the GPe 3 days prior to the recording. Slices of the GPe were examined for accuracy in the injection location. Animals with incorrect injection locations were not used for recording.

### Data analysis and statistics

All statistics are indicated where used. Statistical analyses were performed with GraphPad Prism Software (GraphPad Software, Inc., La Jolla, CA). Normality was tested by D’Agostino-Pearson or Shapiro-Wilk normality tests. Non-normal datasets were log-transformed for normality before statistical testing. All behavioral experiments were controlled by computer systems, and data were collected and analyzed in an automated and unbiased way. Virus-injected animals in which the injection site was incorrect were excluded. No other animals were excluded.

## Results

### Female-specific increase in fear generalization in *16p11.2 del*/+ mice

One hallmark of anxiety disorders is fear generalization (Dunsmoor & Paz, 2015). Fear generalization can be assessed in mice using a fear conditioning paradigm with a discrimination component (see Methods), in which mice are trained to associate one auditory stimulus (conditioned stimulus, CS) (CS+) with a foot shock (unconditioned stimulus, US), while a different auditory stimulus (CS–) is presented without the shock. In a fear retrieval test 24 hours following the conditioning, both freezing in response to the CS+ and that to the CS– are measured and used to calculate a discrimination index, which is the difference between an animal’s average freezing to the CS+ and that to the CS–, normalized to the sum of the two measurements.

Interestingly, we found that during a habituation session before the conditioning, female *16p11.2 del*/+ mice showed small (10-20%) but robust increase in freezing to the auditory stimuli compared with their wildtype (WT) littermates (Figure 1A, left). Male *16p11.2* mice did not show such change (Figure 1A, right). However, we did not observe a significant difference in freezing during the first tone presentation in the subsequent conditioning session (i.e., before mice received any shocks) between genotypes for either the female or the male mice (Figure 1B, D), suggesting that the enhanced freezing in *16p11.2 del*/+ female mice during habituation may be related to the fact that the auditory stimuli were novel to the animals.

**Figure 1.**
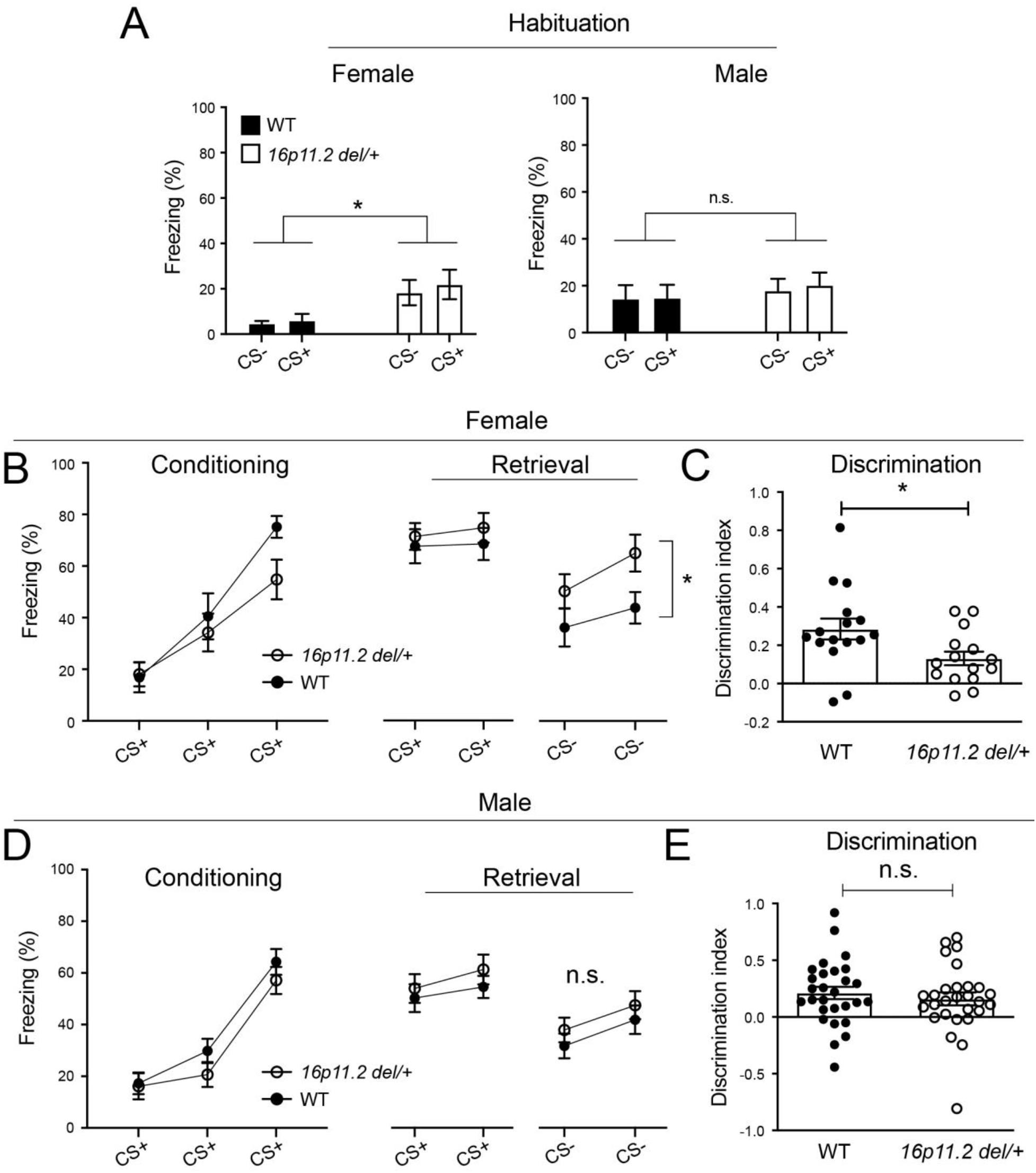
Female *16p11.2 del*/+ mice exhibit fear generalization following fear conditioning. (A) Freezing behavior of male and female *16p11.2 del*/+ mice and their respective wildtype (WT) littermates in response to CS+ and CS– during habituation (female (*16p11.2 del*/+, n = 15 mice, WT, n = 16), F(1, 29) = 6.023, P = 0.0204; male (*16p11.2 del*/+, n = 28 mice, WT, n = 28), F(1, 54) = 0.3433, P = 0.5604; *P < 0.05, n.s., nonsignificant; two-way ANOVA with repeated measures). (B) Freezing to each stimulus presentation during conditioning and retrieval for female mice (conditioning, F(1,29) = 1.419, P = 0.2432; CS+ retrieval, F(1,29) = 0.4314, P = 0.5165; CS–retrieval, F(1,29) = 5.765, P = 0.0230; *p < 0.05; two-way ANOVA with repeated measures and post-hoc Sidak’s test). (C) Discrimination index [(CS^+^ – CS^-^) / (CS^+^ + CS^-^)] for female mice (*P = 0.0192, Mann-Whitney t-test,). (D) Freezing to each stimulus presentation during conditioning and retrieval for male mice. (conditioning, F(1,54) = 0.9938, P = 0.3233; CS+ retrieval, F(1,54) = 0.6327, P = 0.4298; CS–retrieval, F(1,54) = 0.8779, P = 0.3530; two-way ANOVA with repeated measures). (E) Discrimination index for male mice (P = 0.3742, n.s., nonsignificant, Mann-Whitney t-test). Data are presented as mean ± s.e.m.

After fear conditioning and upon memory retrieval, both female and male *16p11.2 del*/+ mice showed levels of freezing similar to those of their WT littermates in response to the CS+ (Figure B, D), consistent with previous findings that *16p11.2 del*/+ mice have intact fear conditioning (Lynch et al., 2020; Yang, Lewis, et al., 2015). Surprisingly, however, female, but not male *16p11.2 del*/+ mice showed increased freezing to the CS– compared with WT littermates (Figure 1B, D), resulting in reduced levels of fear discrimination in female, but not male *16p11.2 del*/+ animals (Figure 1C, E). In addition, we found that female, but not male *16p11.2 del*/+ mice showed enhanced reactions to foot shocks compared with WT mice, as measured by enhanced movement velocity and distance immediately following shocks of varying intensities (Figure 2). These results suggest that female *16p11.2 del*/+ mice have enhanced fear generalization following fear conditioning, which could result from heightened alertness (as indicated by increased freezing during habituation) or an increase in sensitivity to aversive stimuli (as indicated by increase reactivity to foot shocks), or both.

**Figure 2.**
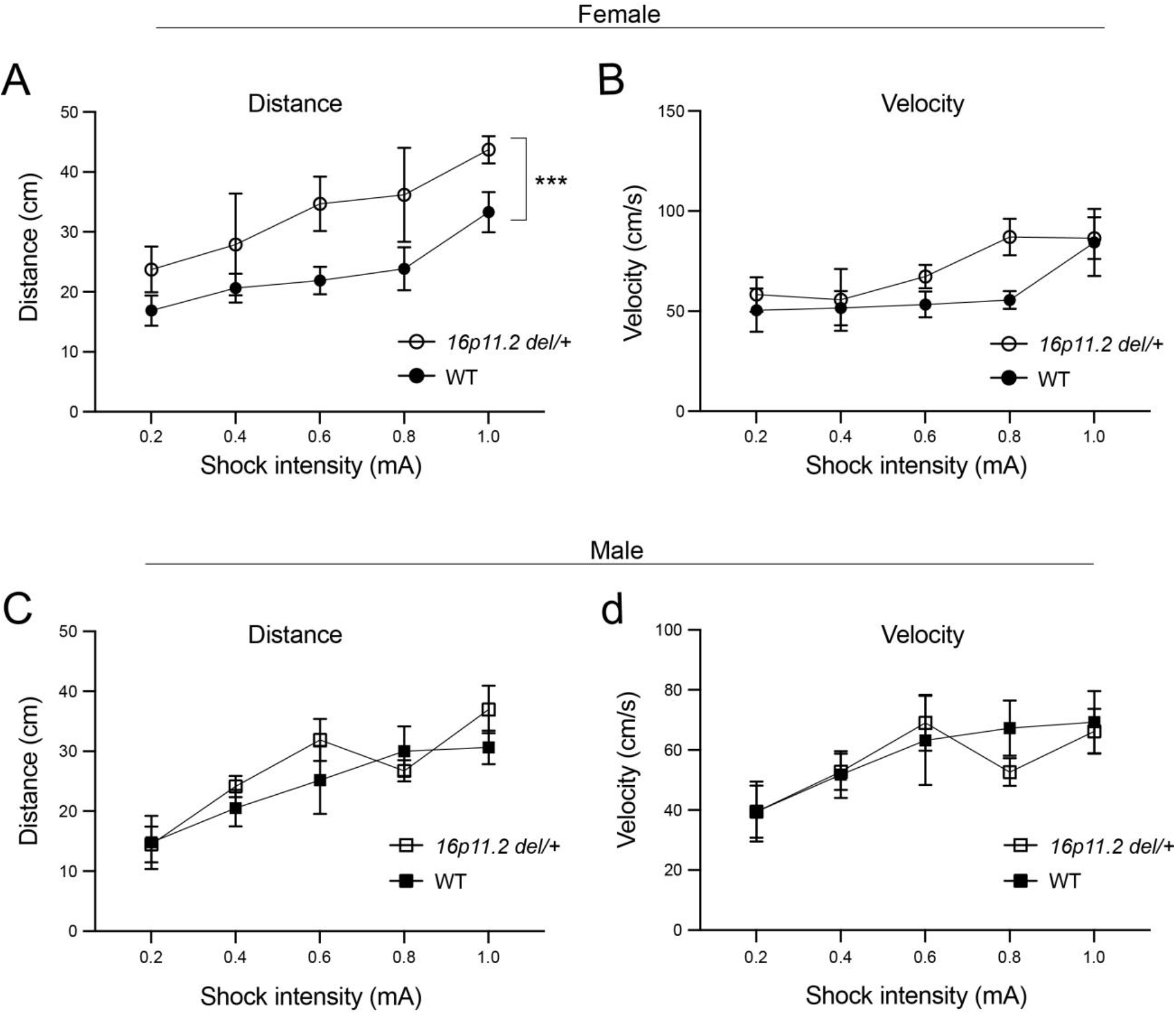
Female *16p11.2 del*/+ mice show enhanced reactivity to foot shock. (A) Distance traveled during 2-s shock presentations for female mice (F(1,50) = 14.94, P = 0.0003; ***P < 0.001; two-way ANOVA; *16p11.2 del*/+, n = 4; WT, n = 8). (B) Movement velocity during 2-s shock presentations for female mice (F(1,50) = 2.596, P = 0.1135; two-way ANOVA). (C) Distance traveled during 2-s shock presentations for male mice (F(1,50) = 1.410, P = 0.2407; two-way ANOVA; *16p11.2 del*/+, n = 7; WT, n = 5). (D) Movement velocity during 2-s shock presentations for male mice (F(1,50) = 0.1467, P = 0.7033; two-way ANOVA). Data are presented as mean ± s.e.m.

### *16p11.2 del*/+ mice have normal auditory perception

An alternative explanation for the enhanced fear generalization in female *16p11.2 del*/+ mice is that these mice have an impairment in auditory processing, such that they cannot effectively discriminate between a 4-kHz tone and a 12-kHz tone, which were used as CS+ and CS–, respectively, during fear conditioning. To test this possibility, we trained a new cohort of mice, including *16p11.2 del*/+ mice and their WT littermates, in an auditory two-alternative choice (2-AC) task that depended on discriminating between a 4-kHz tone and a 12-kHz tone (Figure 3A; see Methods) (Ahrens et al., 2015). Both female and male *16p11.2 del*/+ mice learned the 2-AC task at a rate similar to that of their WT littermates (Figure 3B, C). In fact, male *16p11.2 del*/+ mice tended to be faster than WT mice in learning the task (Figure 3C), though this difference did not reach significance. In addition, the performance of female and male *16p11.2 del*/+ mice in discriminating a series of sounds with frequencies ranging from 4 to 12 kHz (Figure 3D-F and H-J), or with different intensities (Figure 3G, K), was indistinguishable from their WT littermates. These results indicate that *16p11.2* microdeletion does not affect auditory perception in mice, ruling out the possibility that the enhanced fear generalization in female *16p11.2 del*/+ mice is confounded by an impairment in auditory processing in these mice.

**Figure 3.**
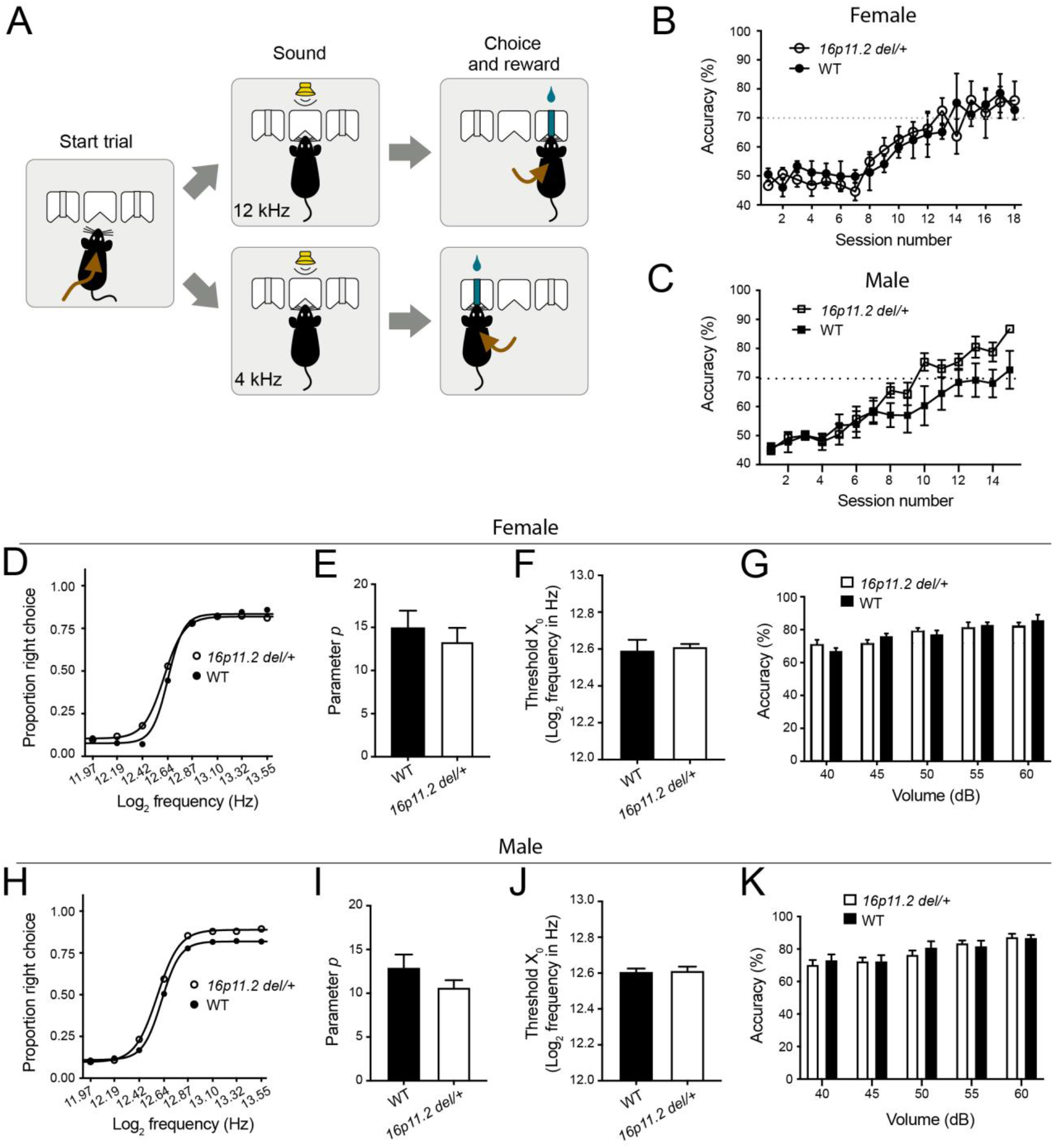
*16p11.2 del*/+ mice have normal auditory perception. (A) A schematic of the behavioral task. (B) Performance levels across training for female mice (F(1,8) = 0.005112, P = 0.9448; two-way ANOVA; *16p11.2 del*/+, n = 7, WT, n = 3). (C) Performance levels across training for male mice (F(1,14) = 2.557, P =0.1321; two-way ANOVA; *16p11.2 del*/+, n = 9; WT, n = 7). (D) Psychometric response curve for frequencies between 4 and 12 kHz (in Log2 values) for female mice. (E) Quantification of the slope of the psychometric curve (parameter *p*) for female mice (P = 0.5878, t-test). (F) Quantification of the median threshold, *Xo*, from the psychometric function for female mice (P = 0.6465, t-test). (G) Average performance levels at 4 and 12 kHz for stimuli volume between 40 and 60 dB for female mice (F(1,8) = 0.04474, P = 0.8378; two-way ANOVA with repeated measures). (H) Psychometric response curve for frequencies between 4 and 12 kHz (in Log2 values) for male mice. (I) Quantification of the slope of the psychometric curve (parameter *p*) for male mice (P = 0.1713, t-test,). (J) Quantification of the median threshold, *Xo*, from the psychometric function for male mice (P = 0.8607, t-test). (K) Average performance levels at 4 and 12 kHz for stimuli volume between 40 and 60 dB for male mice (F(1,14) = 0.0173, P = 0.8972; two-way ANOVA with repeated measures). All data are presented as mean ± s.e.m.

### Stress induces an increase in anxiety in female *16p11.2 del*/+ mice

In fear conditioning, mice receive electric shocks as the aversive US, which is a significant stressor to animals. Therefore, the enhanced fear generalization in female *16p11.2 del*/+ mice after fear conditioning points to a possibility that these animals are prone to stress-induced anxiety. To further test this possibility, we sought to examine anxiety-like behaviors in mice subjected to a different stressor. For this purpose, we used physical restraint (Methods), a well characterized stress-induction procedure in rodents which has been shown to affect the function of the central amygdala (Varodayan et al., 2018, 2019). As described previously (Zimprich et al., 2014), animals are physically restrained in a well-ventilated 50 mL conical tube for 2 hours in a dark, sound attenuated box. 24 hours later, animals were tested on the EPM (Methods). We found a significant interaction between sex and genotype in the time spent in the open arms (Figure 4A) and significant effects of sex on movement velocity (Figure 4B) and distance traveled (Figure 4C). Post-hoc analysis revealed that the stressed female *16p11.2 del*/+ mice spent significantly less time in the open arms of the EPM compared to their female WT littermates (Figure 4A). We did not find any change in time spent in the open arms in male *a16p11.2 del*/+ mice.

**Figure 4.**
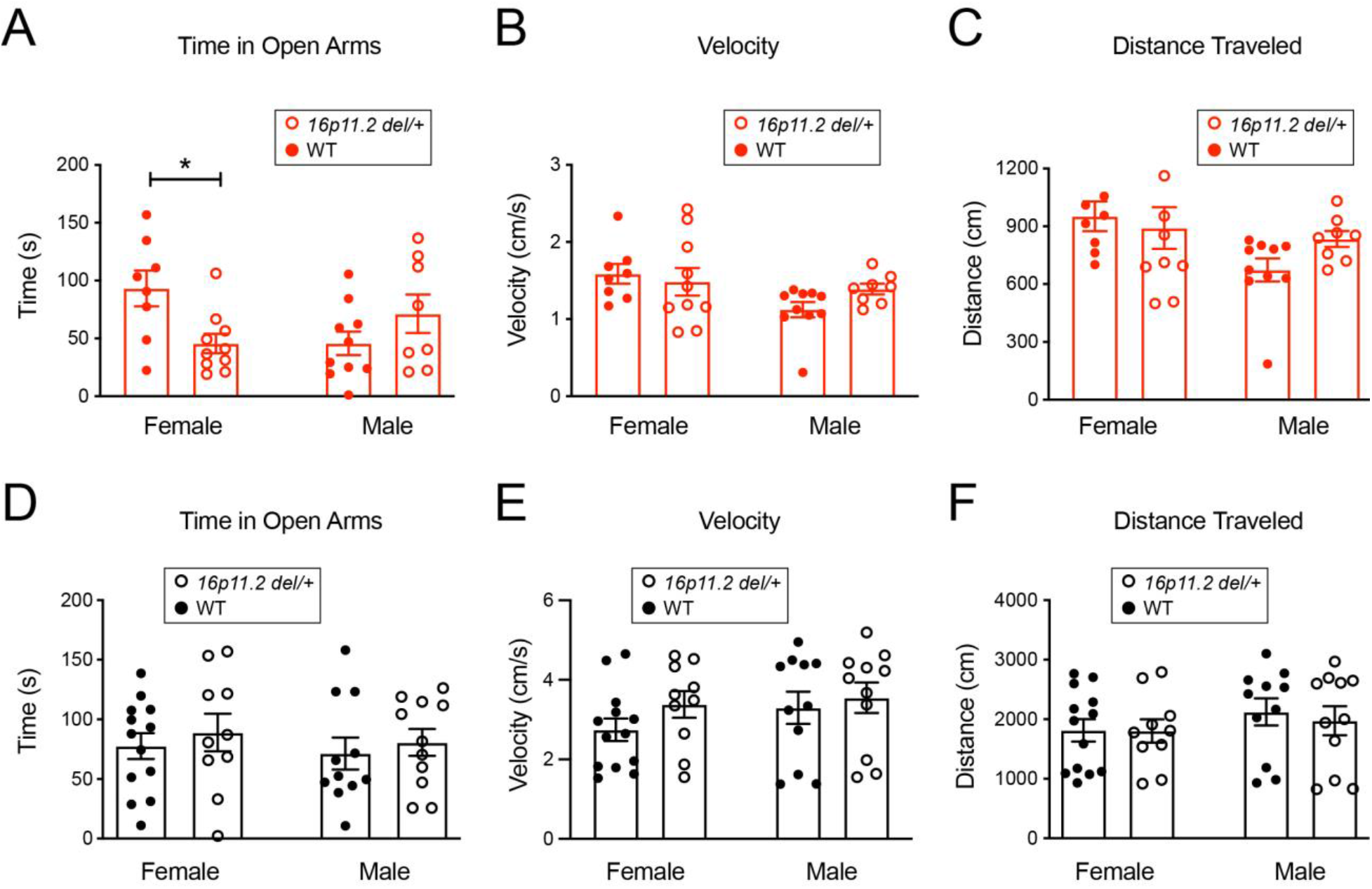
Female *16p11.2 del*/+ mice exhibit enhanced stress-induced anxiety-like behavior. (A) Time spent in the open arms of EPM 24 hours after stress exposure (F(1,32) = 8.553, P = 0.0063; *P < 0.05; two-way ANOVA with post-hoc Sidak’s test; female *16p11.2 del*/+, n = 10, female WT, n = 8, male *16p11.2 del*/+, n = 8, male WT, n = 10). (B) Movement velocity on the EPM 24 hours after stress exposure (F(1,32) = 0.3917, P = 0.5358; two-way ANOVA). Same mice as in A are used. (C) Distance traveled on the EPM 24 hours after stress exposure (F(1,32) = 0.3918, P = 0.5358; two-way ANOVA). Same mice as in A are used. (D) Time spent in the open arms of EPM in naïve mice (F(1,41) = 0.6545, P = 0.4232; two-way ANOVA; female *16p11.2 del*/+, n = 10, female WT, n = 13, male *16p11.2 del*/+, n = 11; male WT, n = 11). (E) Movement velocity on the EPM in naïve mice (F(1,41) = 1.587, P = 0.2148; two-way ANOVA). Same mice as in D are used. (F) Distance traveled on the EPM in naïve mice (F(1,41) = 0.1314 P = 0.7189; two-way ANOVA). Same mice as in D are used. Data are presented as mean ± s.e.m.

We also examined anxiety levels in naïve mice using the EPM test. Compared with naïve female or male WT littermates, naïve female or male *16p11.2 del*/+ mice, respectively, did not show any change in the time spent in the open arms (Figure 4D), movement velocity (Figure 4E) and distance traveled (Figure 4F). This result is consistent with previous findings (Arbogast et al., 2016; Lynch et al., 2020; Yang, Lewis, et al., 2015). Together, our results indicate that female *16p11.2 del*/+ mice have increased susceptibility to stress-induced anxiety.

### *16p11.2 del*/+ mice have central amygdala dysfunction

Previous studies have revealed that the central amygdala (CeA) is particularly responsive to stress and is a major contributor to anxiety-related behaviors (Ahrens et al., 2018; Fox et al., 2012; Shackman & Fox, 2016). Therefore, we examined whether the *16p11.2* microdeletion affects CeA neuronal function in a sex-specific manner. We recorded miniature excitatory postsynaptic currents (mEPSCs) – a measurement of total excitatory synaptic drive onto the recorded neurons – from CeA neurons in acute brain slices prepared from female or male *16p11.2 del*/+ mice, as well as their respective WT littermates (Figure 5A). We found significant effects of sex and genotype on mEPSC frequency in randomly recorded central amygdala neurons (Figure 5B-D). Post-hoc analysis revealed that females with 16p11.2 microdeletion specifically had increased mEPSC frequency compared with female wildtype littermates. There was no difference in mEPSC amplitude between genotypes or sexes (Figure 5E). These results indicate that female, but not male *16p11.2 del*/+ mice have enhanced excitatory synaptic drive onto CeA neurons.

**Figure 5.**
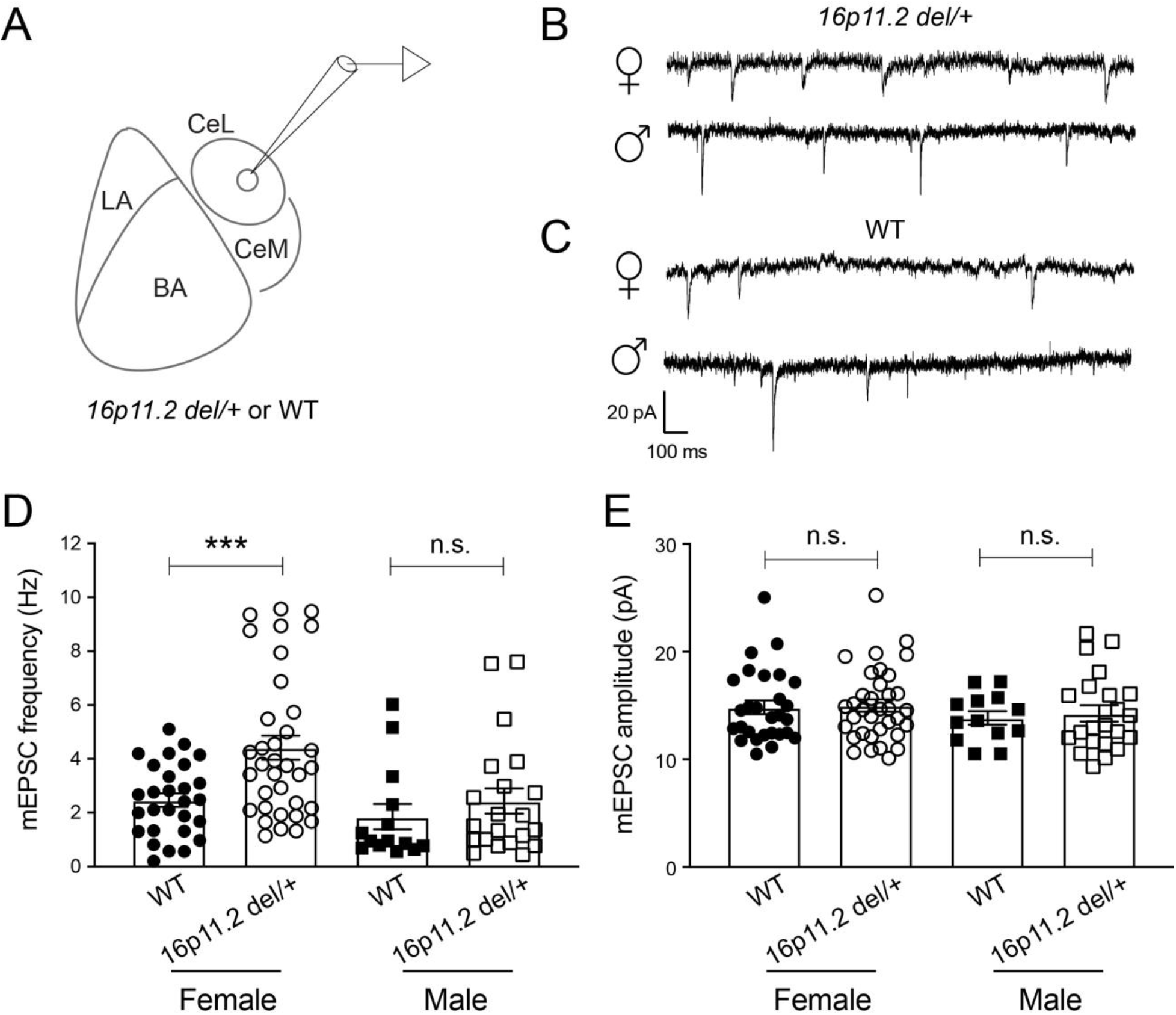
Female *16p11.2 del*/+ mice have increased excitatory synaptic transmission onto CeA neurons. (A) A schematic of the experimental design. (B, C) Representative mEPSC traces from CeA neurons recorded from male and female *16p11.2 del*/+ (B) and WT (C) mice. (D) Quantification of mEPSC frequency for CeA neurons (F(1, 94) = 7.759, P = 0.0065; ***P < 0.001; two-way ANOVA with post-hoc Sidak’s test; female *16p11.2 del*/+, n = 35 cells from 4 mice, female WT, n = 28 cells from 3 mice, male *16p11.2 del*/+, n = 21 cells from 4 mice, male WT, n = 14 cells from 3 mice). (E) Quantification of mEPSC amplitude for CeA neurons (F(1,94) = 0.1620, P = 0.6882; two-way ANOVA). Data are from the same cells as in D. Data are presented as mean ± s.e.m.

We recently identified a pathway from the CeA to the globus pallidus externa (GPe), which conveys information of the US and is critical for learning in fear conditioning (Giovanniello et al., 2020). Importantly, optogenetic activation of the CeA-GPe pathway increases fear generalization whereby animals increase their freezing to CS–. Therefore, we sought to determine whether the GPe-projecting CeA neurons are affected by the 16p11.2 microdeletion. To this end, we used a retrograde labeling strategy whereby fluorescently conjugated CTB was injected in the GPe to label the GPe-projecting CeA neurons (Figure 6A; Methods). Three days after the CTB injection, we recorded mEPSCs selectively from the CTB-labeled GPe-projecting CeA neurons in acute brain slices prepared from female or male *16p11.2 del*/+ mice, as well as their respective WT littermates (Figure 6A, B). Again, we found a significant interaction between sex and genotype whereby females with 16p11.2 microdeletion exhibited increased mEPSC frequency compared with wildtype littermates (Figure 6D, E). Thus, our results indicate that the *16p11.2* microdeletion caused a female-specific enhancement of excitatory synaptic drive onto CeA neurons, and moreover suggests dysfunction in the CeA-GPe pathway as a potential mechanism for the increased stress susceptibility and fear generalization identified in female *16p11.2 del*/+ mice.

**Figure 6.**
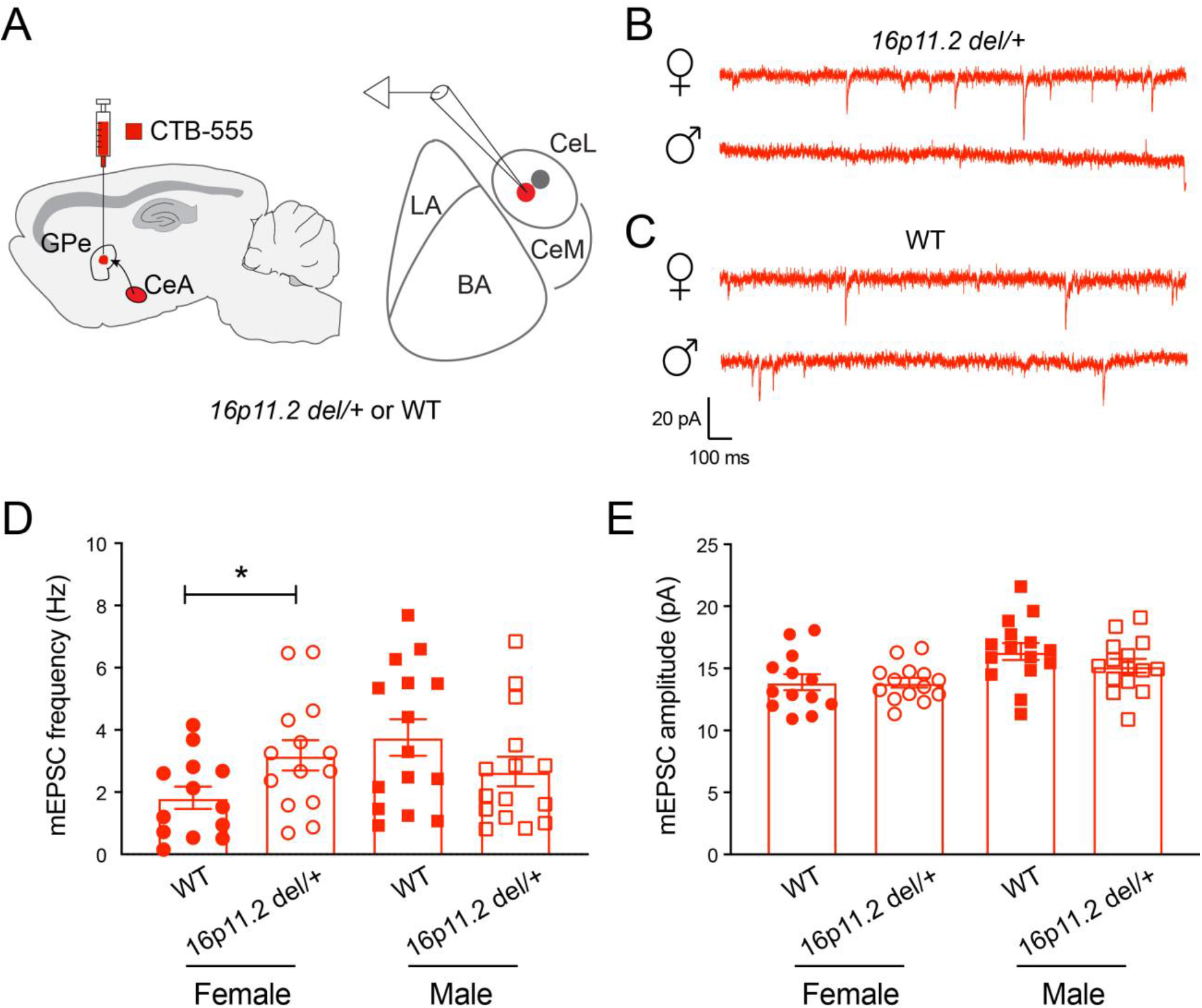
Female *16p11.2 del*/+ mice have increased excitatory synaptic transmission onto GPe-projecting CeA neurons. (A) A schematic of the experimental design. CTB-555 was used to retrogradely label GPe-projecting CeA neurons. (B, C) Representative mEPSC traces from GPe-projecting CeA neurons recorded from male and female *16p11.2 del*/+ (B) and WT (C) mice. (D) Quantification of mEPSC frequency for GPe-projecting CeA neurons (F(1, 53) = 6.251, P = 0.0155; *P < 0.05; two-way ANOVA with post-hoc Sidak’s test; female *16p11.2 del*/+, n = 14 cells from 5 mice, female WT, n = 13 cells from 7 mice, male *16p11.2 del*/+, n = 15 cells from 3 mice, male WT, n = 15 cells from 5 mice). (E) Quantification of mEPSC amplitude for GPe-projecting CeA neurons (F(1,53) = 1.055, P = 0.3090; two-way ANOVA). Data are from the same cells as in D. Data are presented as mean ± s.e.m.

## Discussion

Our results indicate that female, but not male, *16p11.2 del*/+ mice have increased susceptibility to anxiety-like phenotypes following stressful life events, revealing a previously underappreciated sex-specific effect in the modulation of behavior by 16p11.2 microdeletion. Furthermore, we identify that CeA dysfunction, in particular that in the CeA-GPe circuit, may underlie the female-specific behavioral phenotypes caused by the 16p11.2 microdeletion. These findings are consistent with the vast literature that females affected with ASD show distinct behavioral symptoms compared with males (Beggiato et al., 2017; Dean et al., 2017; Head et al., 2014; Hiller et al., 2014; Werling & Geschwind, 2013) in particular the more internalizing symptoms and emotional disturbances (Horiuchi et al., 2014; Kreiser & White, 2014; Rynkiewicz et al., 2016; Rynkiewicz & Łucka, 2018; Solomon et al., 2012). Our findings are also consistent with the notion that in the general population, females have an increased prevalence of stress-related disorders such as anxiety, depression, and PTSD (Breslau, 2002; Kessler et al., 1995; Olff, 2017; Tolin & Foa, 2006). Our study thus urges a careful examination of anxiety and other emotional symptoms, as well as functional changes in the amygdala-basal ganglia circuits in 16p11.2 microdeletion carriers, in particular in female carriers. In general, our study also urges sex-specific diagnostic and treatment strategies for ASD.

Three lines of evidence suggest that heightened alertness or an increase in sensitivity to aversive stimuli, or to the stimuli signaling potential threat, may underlie the increased susceptibility to anxiety-like phenotypes in female *16p11.2 del*/+ mice following stressful experiences. First, *16p11.2 del*/+ mice, especially females, show increased freezing when they are exposed to an unfamiliar sound, which is a sign of uncertainty or potential danger. Second, female *16p11.2 del*/+ mice have enhanced reactivity to foot shocks. Third, CeA neurons in female *16p11.2 del*/+ mice have enhanced sensitivity to excitatory inputs. This enhanced sensitivity may lead to heightened alertness or attention, as the CeA has been implicated in selective processing of salient information (Calu et al., 2010; Roesch et al., 2012).

The CeA has central roles in the generation of fear and anxiety states (Ahrens et al., 2018; Andreatta et al., 2015; Calhoon & Tye, 2015; Davis et al., 2010; Etkin & Wager, 2007; Fox et al., 2012, 2015; Gungor & Paré, 2016; Jennings et al., 2013; S.-Y. Kim et al., 2013; Li et al., 2013; Marcinkiewcz et al., 2016; Mobbs et al., 2010; Penzo et al., 2015; Shackman & Fox, 2016; Tovote et al., 2015; Wager et al., 2008; Walker & Davis, 2008; Yu et al., 2017). In parallel, amygdala dysfunction has been implicated in the pathogenesis of ASD. Abnormal developmental trajectory of the amygdala has been observed in ASD (Amaral et al., 2008). Brain imaging studies indicate that the amygdala is enlarged precociously in children with autism (Schumann et al., 2004; Sparks et al., 2002), and that amygdala enlargement in autistic children is associated with anxiety symptoms (Juranek et al., 2006). In addition, cellular changes in the amygdala have been reported in ASD (Amaral et al., 2008). In a recent study (Giovanniello et al., 2020), we found that a subpopulation of neurons in the CeA send direct projections to the GPe, and the CeA-GPe pathway conveys US information and controls learning during fear conditioning. In the current study, we found that an enhanced excitatory drive onto GPe-projecting CeA neurons parallels the anxiety phenotypes of female *16p11.2 del*/+ mice. These findings together strongly suggest a role of CeA-GPe circuit dysfunction in susceptibility to anxiety after stress, and warrant future studies to elucidate how this circuit is involved in 16p11.2 microdeletion syndrome.

## Acknowledgements

We thank members of the Li laboratory for helpful discussions, Pavel Osten for providing *16p11.2 Del*/+ breeders, and Alea Mills for providing primers for genotyping the *16p11.2 del*/+ mice. This work was supported by grants from the National Institutes of Health (NIH) (R01MH101214, R21MH114070, B.L.), Simons Foundation (344904, B.L.), National Alliance for Research on Schizophrenia and Depression (Grant 23169, B.L.; Grant 21227, S.A.), the Cold Spring Harbor Laboratory and Northwell Health Affiliation (B.L.) and Feil Family Neuroscience Endowment (B.L.).

## Author contributions

J.G. and B.L. conceived and designed the study. J.G. conducted the experiments and analyzed data. S.A. conducted the experiments with the auditory discrimination task and assisted with other electrophysiology experiments. K.Y. identified the CeA-GPe projections and assisted with experiments. J.G. and B.L. wrote the paper with input from all authors.

## References

Ahrens, S., Jaramillo, S., Yu, K., Ghosh, S., Hwang, G.-R., Paik, R., Lai, C., He, M., Huang, Z. J., & Li, B. (2015). ErbB4 regulation of a thalamic reticular nucleus circuit for sensory selection. Nature Neuroscience, 18(1), 104–111. https://doi.org/10.1038/nn.3897

Ahrens, S., Wu, M. V., Furlan, A., Hwang, G.-R., Paik, R., Li, H., Penzo, M. A., Tollkuhn, J., & Li, B. (2018). A Central Extended Amygdala Circuit That Modulates Anxiety. The Journal of Neuroscience, 38(24), 5567–5583. https://doi.org/10.1523/jneurosci.0705-18.2018

Al-Kateb, H., Khanna, G., Filges, I., Hauser, N., Grange, D. K., Shen, J., Smyser, C. D., Kulkarni, S., & Shinawi, M. (2014). Scoliosis and vertebral anomalies: Additional abnormal phenotypes associated with chromosome 16p11.2 rearrangement. American Journal of Medical Genetics Part A, 164(5), 1118–1126. https://doi.org/10.1002/ajmg.a.36401

Amaral, D. G., Schumann, C. M., & Nordahl, C. W. (2008). Neuroanatomy of autism. Trends in Neurosciences, 31(3), 137–145. https://doi.org/10.1016/j.tins.2007.12.005

Andersen, P. N., Skogli, E. W., Hovik, K. T., Egeland, J., & Øie, M. (2015). Associations Among Symptoms of Autism, Symptoms of Depression and Executive Functions in Children with High-Functioning Autism: A 2 Year Follow-Up Study. Journal of Autism and Developmental Disorders, 45(8), 2497–2507. https://doi.org/10.1007/s10803-015-2415-8

Andreatta, M., Glotzbach-Schoon, E., Mühlberger, A., Schulz, S. M., Wiemer, J., & Pauli, P. (2015). Initial and sustained brain responses to contextual conditioned anxiety in humans. Cortex, 63, 352–363. https://doi.org/10.1016/j.cortex.2014.09.014

Angelakos, C. C., Watson, A. J., O’Brien, W. T., Krainock, K. S., Nickl-Jockschat, T., & Abel, T. (2017). Hyperactivity and male-specific sleep deficits in the 16p11.2 deletion mouse model of autism. Autism Research, 10(4), 572–584. https://doi.org/10.1002/aur.1707

Antshel, K. M., & Russo, N. (2019). Autism Spectrum Disorders and ADHD: Overlapping Phenomenology, Diagnostic Issues, and Treatment Considerations. Current Psychiatry Reports, 21(5), 34. https://doi.org/10.1007/s11920-019-1020-5

Antshel, K. M., Zhang-James, Y., & Faraone, S. V. (2014). The comorbidity of ADHD and autism spectrum disorder. Expert Review of Neurotherapeutics, 13(10), 1117–1128. https://doi.org/10.1586/14737175.2013.840417

Antshel, K. M., Zhang-James, Y., Wagner, K. E., Ledesma, A., & Faraone, S. V. (2016). An update on the comorbidity of ADHD and ASD: a focus on clinical management. Expert Review of Neurotherapeutics, 16(3), 1–15. https://doi.org/10.1586/14737175.2016.1146591

Arbogast, T., Ouagazzal, A.-M., Chevalier, C., Kopanitsa, M., Afinowi, N., Migliavacca, E., Cowling, B. S., Birling, M.-C., Champy, M.-F., Reymond, A., & Herault, Y. (2016). Reciprocal Effects on Neurocognitive and Metabolic Phenotypes in Mouse Models of 16p11.2 Deletion and Duplication Syndromes. PLOS Genetics, 12(2), e1005709. https://doi.org/10.1371/journal.pgen.1005709

Baio, J., Wiggins, L., Christensen, D. L., Maenner, M. J., Daniels, J., Warren, Z., Kurzius-Spencer, M., Zahorodny, W., Robinson, C., Rosenberg, White, T., Durkin, M. S., Imm, P., Nikolaou, L., Yeargin-Allsopp, M., Lee, L.-C., Harrington, R., Lopez, M., Fitzgerald, R. T.,… Dowling, N. F. (2018). Prevalence of Autism Spectrum Disorder Among Children Aged 8 Years — Autism and Developmental Disabilities Monitoring Network, 11 Sites, United States, 2014. MMWR Surveillance Summaries, 67(6), 1–23. https://doi.org/10.15585/mmwr.ss6706a1

Baron-Cohen, S. (2009). Autism: The Empathizing–Systemizing (E-S) Theory. Annals of the New York Academy of Sciences, 1156(1), 68–80. https://doi.org/10.1111/j.1749-6632.2009.04467.x

Beery, A. K., & Zucker, I. (2011). Sex bias in neuroscience and biomedical research. Neuroscience & Biobehavioral Reviews, 55(3), 565–572. https://doi.org/10.1016/j.neubiorev.2010.07.002

Beggiato, A., Peyre, H., Maruani, A., Scheid, I., Rastam, M., Amsellem, F., Gillberg, C. I., Leboyer, M., Bourgeron, T., Gillberg, C., & Delorme, R. (2017). Gender differences in autism spectrum disorders: Divergence among specific core symptoms. Autism Research, 10(4), 680–689. https://doi.org/10.1002/aur.1715

Bijlsma, E. K., Gijsbers, A. C. J., Schuurs-Hoeijmakers, J. H. M., Haeringen, A. van, Putte, D. E. F. van de, Anderlid, B.-M., Lundin, J., Lapunzina, P., Jurado, L. A. P., Chiaie, B. D., Loeys, B., Menten, B., Oostra, A., Verhelst, H., Amor, D. J., Bruno, D. L., Essen, A. J. van, Hordijk, R., Sikkema-Raddatz, B.,… Ruivenkamp, C. A. L. (2009). Extending the phenotype of recurrent rearrangements of 16p11.2: Deletions in mentally retarded patients without autism and in normal individuals. European Journal of Medical Genetics, 52(2–3), 77–87. https://doi.org/10.1016/j.ejmg.2009.03.006

Bölte, S., Duketis, E., Poustka, F., & Holtmann, M. (2011). Sex differences in cognitive domains and their clinical correlates in higher-functioning autism spectrum disorders. Autism, 15(4), 497–511. https://doi.org/10.1177/1362361310391116

Breslau, N. (2002). Gender differences in trauma and posttraumatic stress disorder. The Journal of Gender-Specific Medicine: JGSM: The Official Journal of the Partnership for Women’s Health at Columbia, 5(1), 34–40.

Brookman-Frazee, L., Stadnick, N., Chlebowski, C., Baker-Ericzén, M., & Ganger, W. (2018). Characterizing psychiatric comorbidity in children with autism spectrum disorder receiving publicly funded mental health services. Autism, 22(8), 938–952. https://doi.org/10.1177/1362361317712650

Calhoon, G. G., & Tye, K. M. (2015). Resolving the neural circuits of anxiety. Nature Neuroscience, 18(10), 1394–1404. https://doi.org/10.1038/nn.4101

Calu, D. J., Roesch, M. R., Haney, R. Z., Holland, P. C., & Schoenbaum, G. (2010). Neural Correlates of Variations in Event Processing during Learning in Central Nucleus of Amygdala. Neuron, 68(5), 991–1001. https://doi.org/10.1016/j.neuron.2010.11.019

Chen, C.-H., Chen, H.-I., Chien, W.-H., Li, L.-H., Wu, Y.-Y., Chiu, Y.-N., Tsai, W.-C., & Gau, S. S.-F. (2017). High resolution analysis of rare copy number variants in patients with autism spectrum disorder from Taiwan. Scientific Reports, 7(1), 11919. https://doi.org/10.1038/s41598-017-12081-4

Dalla, C., & Shors, T. J. (2009). Sex differences in learning processes of classical and operant conditioning. Physiology & Behavior, 97(2), 229–238. https://doi.org/10.1016/j.physbeh.2009.02.035

Davidsson, M., Hult, N., Gillberg, C., Särneö, C., Gillberg, C., & Billstedt, E. (2017). Anxiety and depression in adolescents with ADHD and autism spectrum disorders; correlation between parent-and self-reports and with attention and adaptive functioning. Nordic Journal of Psychiatry, 71(8), 1–7. https://doi.org/10.1080/08039488.2017.1367840

Davis, M., Walker, D. L., Miles, L., & Grillon, C. (2010). Phasic vs Sustained Fear in Rats and Humans: Role of the Extended Amygdala in Fear vs Anxiety. Neuropsychopharmacology, 35(1), 105–135. https://doi.org/10.1038/npp.2009.109

Dean, M., Harwood, R., & Kasari, C. (2017). The art of camouflage: Gender differences in the social behaviors of girls and boys with autism spectrum disorder. Autism, 21(6), 678–689. https://doi.org/10.1177/1362361316671845

Diagnostic and Statistical Manual of Mental Disorders, DSM-5 (5th ed.). (2013). American Psychiatric Association. https://doi.org/10.5555/appi.books.9780890425596.x00pre

Dunsmoor, J. E., & Paz, R. (2015). Fear Generalization and Anxiety: Behavioral and Neural Mechanisms. Biological Psychiatry, 78(5), 336–343. https://doi.org/10.1016/j.biopsych.2015.04.010

Etkin, A., & Wager, T. D. (2007). Functional Neuroimaging of Anxiety: A Meta-Analysis of Emotional Processing in PTSD, Social Anxiety Disorder, and Specific Phobia. American Journal of Psychiatry, 164(10), 1476–1488. https://doi.org/10.1176/appi.ajp.2007.07030504

Fernandez, B. A., Roberts, W., Chung, B., Weksberg, R., Meyn, S., Szatmari, P., Joseph-George, A. M., MacKay, S., Whitten, K., Noble, B., Vardy, C., Crosbie, V., Luscombe, S., Tucker, E., Turner, L., Marshall, C. R., & Scherer, S. W. (2010). Phenotypic spectrum associated with de novo and inherited deletions and duplications at 16p11.2 in individuals ascertained for diagnosis of autism spectrum disorder. Journal of Medical Genetics, 47(3), 195. https://doi.org/10.1136/jmg.2009.069369

Fombonne, E. (2002). Epidemiological trends in rates of autism. Molecular Psychiatry, 7(S2), S4. https://doi.org/10.1038/sj.mp.4001162

Fox, A. S., Oler, J. A., Shelton, S. E., Nanda, S. A., Davidson, R. J., Roseboom, P. H., & Kalin, N. H. (2012). Central amygdala nucleus (Ce) gene expression linked to increased trait-like Ce metabolism and anxious temperament in young primates. Proceedings of the National Academy of Sciences, 109(44), 18108–18113. https://doi.org/10.1073/pnas.1206723109

Fox, A. S., Oler, J. A., Tromp, D. P. M., Fudge, J. L., & Kalin, N. H. (2015). Extending the amygdala in theories of threat processing. Trends in Neurosciences, 38(5), 319–329. https://doi.org/10.1016/j.tins.2015.03.002

Giarelli, E., Wiggins, L. D., Rice, C. E., Levy, S. E., Kirby, R. S., Pinto-Martin, J., & Mandell, D. (2010). Sex differences in the evaluation and diagnosis of autism spectrum disorders among children. Disability and Health Journal, 3(2), 107–116. https://doi.org/10.1016/j.dhjo.2009.07.001

Gilchrist, J. M., Jerwood, D., & Ismaiel, H. S. (2005). Comparing and unifying slope estimates across psychometric function models. Perception & Psychophysics, 67(7), 1289–1303. https://doi.org/10.3758/bf03193560

Giovanniello, J., Yu, K., Furlan, A., Nachtrab, G. T., Sharma, R., Chen, X., & Li, B. (2020). A central amygdala-globus pallidus circuit conveys unconditioned stimulus-related information and controls fear learning. The Journal of Neuroscience, JN-RM-2090-20. https://doi.org/10.1523/jneurosci.2090-20.2020

Gotham, K., Bishop, S. L., Hus, V., Huerta, M., Lund, S., Buja, A., Krieger, A., & Lord, C. (2013). Exploring the Relationship Between Anxiety and Insistence on Sameness in Autism Spectrum Disorders. Autism Research, 6(1), 33–41. https://doi.org/10.1002/aur.1263

Grissom, N. M., McKee, S. E., Schoch, H., Bowman, N., Havekes, R., O’Brien, W. T., Mahrt, E., Siegel, S., Commons, K., Portfors, C., Nickl-Jockschat, T., Reyes, T. M., & Abel, T. (2017). Male-specific deficits in natural reward learning in a mouse model of neurodevelopmental disorders. Molecular Psychiatry, 23(3), 544. https://doi.org/10.1038/mp.2017.184

Gruene, T. M., Flick, K., Stefano, A., Shea, S. D., & Shansky, R. M. (2015). Sexually divergent expression of active and passive conditioned fear responses in rats. ELife, 4, e11352. https://doi.org/10.7554/elife.11352

Gungor, N. Z., & Paré, D. (2016). Functional Heterogeneity in the Bed Nucleus of the Stria Terminalis. The Journal of Neuroscience, 36(31), 8038–8049. https://doi.org/10.1523/jneurosci.0856-16.2016

Hartley, S. L., & Sikora, D. M. (2009). Sex Differences in Autism Spectrum Disorder: An Examination of Developmental Functioning, Autistic Symptoms, and Coexisting Behavior Problems in Toddlers. Journal of Autism and Developmental Disorders, 39(12), 1715. https://doi.org/10.1007/s10803-009-0810-8

Head, A. M., McGillivray, J. A., & Stokes, M. A. (2014). Gender differences in emotionality and sociability in children with autism spectrum disorders. Molecular Autism, 5(1), 19. https://doi.org/10.1186/2040-2392-5-19

Hiller, R. M., Young, R. L., & Weber, N. (2014). Sex Differences in Autism Spectrum Disorder based on DSM-5 Criteria: Evidence from Clinician and Teacher Reporting. Journal of Abnormal Child Psychology, 42(8), 1381–1393. https://doi.org/10.1007/s10802-014-9881-x

Hirvikoski, T, Boman, M., Chen, Q., D’Onofrio, B. M., Mittendorfer-Rutz, E., Lichtenstein, P., Bölte, S., & Larsson, H. (2019). Individual risk and familial liability for suicide attempt and suicide in autism: a population-based study. Psychological Medicine, 1–12. https://doi.org/10.1017/s0033291719001405

Hirvikoski, Tatja, Mittendorfer-Rutz, E., Boman, M., Larsson, H., Lichtenstein, P., & Bölte, S. (2016). Premature mortality in autism spectrum disorder. British Journal of Psychiatry, 208(3), 232–238. https://doi.org/10.1192/bjp.bp.114.160192

Horev, G., Ellegood, J., Lerch, J. P., Son, Y.-E. E., Muthuswamy, L., Vogel, H., Krieger, A. M., Buja, A., Henkelman, R. M., Wigler, M., & Mills, A. A. (2011). Dosage-dependent phenotypes in models of 16p11.2 lesions found in autism. Proceedings of the National Academy of Sciences, 108(41), 17076–17081. https://doi.org/10.1073/pnas.1114042108

Horiuchi, F., Oka, Y., Uno, H., Kawabe, K., Okada, F., Saito, I., Tanigawa, T., & Ueno, S. (2014). Age-and sex-related emotional and behavioral problems in children with autism spectrum disorders: Comparison with control children. Psychiatry and Clinical Neurosciences, 68(7), 542–550. https://doi.org/10.1111/pcn.12164

Howlin, P. (2000). Autism and intellectual disability: Diagnostic and treatment issues. Journal of the Royal Society of Medicine, 93(7), 351–355. https://doi.org/10.1177/014107680009300704

Hudac, C. M., Bove, J., Barber, S., Duyzend, M., Wallace, A., Martin, C. L., Ledbetter, D. H., Hanson, E., Goin-Kochel, R. P., Green-Snyder, L., Chung, W. K., Eichler, E. E., & Bernier, R. A. (2020). Evaluating heterogeneity in ASD symptomatology, cognitive ability, and adaptive functioning among 16p11.2 CNV carriers. Autism Research: Official Journal of the International Society for Autism Research. https://doi.org/10.1002/aur.2332

Hughes, R. N. (2007). Sex does matter comments on the prevalence of male-only investigations of drug effects on rodent behaviour. Behavioural Pharmacology, 18(7), 583–589. https://doi.org/10.1097/fbp.0b013e3282eff0e8

Jang, J., Matson, J. L., Williams, L. W., Tureck, K., Goldin, R. L., & Cervantes, P. E. (2013). Rates of comorbid symptoms in children with ASD, ADHD, and comorbid ASD and ADHD. Research in Developmental Disabilities, 34(8), 2369–2378. https://doi.org/10.1016/j.ridd.2013.04.021

Jennings, J. H., Sparta, D. R., Stamatakis, A. M., Ung, R. L., Pleil, K. E., Kash, T. L., & Stuber, G. D. (2013). Distinct extended amygdala circuits for divergent motivational states. Nature, 496(7444), 224–228. https://doi.org/10.1038/nature12041

Juranek, J., Filipek, P. A., Berenji, G. R., Modahl, C., Osann, K., & Spence, M. A. (2006). Association Between Amygdala Volume and Anxiety Level: Magnetic Resonance Imaging (MRI) Study in Autistic Children. Journal of Child Neurology, 21(12), 1051–1058. https://doi.org/10.1177/7010.2006.00237

Kalyva, E. (2009). Comparison of Eating Attitudes between Adolescent Girls with and without Asperger Syndrome: Daughters’ and Mothers’ Reports. Journal of Autism and Developmental Disorders, 39(3), 480–486. https://doi.org/10.1007/s10803-008-0648-5

Kessler, R. C., Sonnega, A., Bromet, E., Hughes, M., & Nelson, C. B. (1995). Posttraumatic Stress Disorder in the National Comorbidity Survey. Archives of General Psychiatry, 52(12), 1048–1060. https://doi.org/10.1001/archpsyc.1995.03950240066012

Kim, K., & Han, P. (2006). Optimization of chronic stress paradigms using anxiety-and depression-like behavioral parameters. Journal of Neuroscience Research, 83(3), 497–507. https://doi.org/10.1002/jnr.20754

Kim, S.-Y., Adhikari, A., Lee, S. Y., Marshel, J. H., Kim, C. K., Mallory, C. S., Lo, M., Pak, S., Mattis, J., Lim, B. K., Malenka, R. C., Warden, M. R., Neve, R., Tye, K. M., & Deisseroth, K. (2013). Diverging neural pathways assemble a behavioural state from separable features in anxiety. Nature, 496(7444), 219–223. https://doi.org/10.1038/nature12018

Kirby, A. V., Bakian, A. V., Zhang, Y., Bilder, D. A., Keeshin, B. R., & Coon, H. (2019). A 20-year study of suicide death in a statewide autism population. Autism Research, 12(4), 658–666. https://doi.org/10.1002/aur.2076

Knickmeyer, R. C., Wheelwright, S., & Baron-Cohen, S. B. (2008). Sex-typical Play: Masculinization/Defeminization in Girls with an Autism Spectrum Condition. Journal of Autism and Developmental Disorders, 38(6), 1028–1035. https://doi.org/10.1007/s10803-007-0475-0

Kopp, S., Beckung, E., & Gillberg, C. (2010). Developmental coordination disorder and other motor control problems in girls with autism spectrum disorder and/or attention-deficit/hyperactivity disorder. Research in Developmental Disabilities, 31(2), 350–361. https://doi.org/10.1016/j.ridd.2009.09.017

Kreiser, N. L., & White, S. W. (2014). ASD in Females: Are We Overstating the Gender Difference in Diagnosis? Clinical Child and Family Psychology Review, 17(1), 67–84. https://doi.org/10.1007/s10567-013-0148-9

Kumar, R. A., KaraMohamed, S., Sudi, J., Conrad, D. F., Brune, C., Badner, J. A., Gilliam, T. C., Nowak, N. J., Cook, E. H., Dobyns, W. B., & Christian, S. L. (2007). Recurrent 16p11.2 microdeletions in autism. Human Molecular Genetics, 17(4), 628–638. https://doi.org/10.1093/hmg/ddm376

Lai, M.-C., Lombardo, M. V., & Baron-Cohen, S. (2014). Autism. The Lancet, 383(9920), 896–910. https://doi.org/10.1016/s0140-6736(13)61539-1

Leyfer, O. T., Folstein, S. E., Bacalman, S., Davis, N. O., Dinh, E., Morgan, J., Tager-Flusberg, H., & Lainhart, J. E. (2006). Comorbid Psychiatric Disorders in Children with Autism: Interview Development and Rates of Disorders. Journal of Autism and Developmental Disorders, 36(7), 849–861. https://doi.org/10.1007/s10803-006-0123-0

Li, H., Penzo, M. A., Taniguchi, H., Kopec, C. D., Huang, Z. J., & Li, B. (2013). Experience-dependent modification of a central amygdala fear circuit. Nature Neuroscience, 16(3), 332. https://doi.org/10.1038/nn.3322

Lynch, J. F., Ferri, S. L., Angelakos, C., Schoch, H., Nickl-Jockschat, T., Gonzalez, A., O’Brien, W. T., & Abel, T. (2020). Comprehensive Behavioral Phenotyping of a 16p11.2 Del Mouse Model for Neurodevelopmental Disorders. Autism Research. https://doi.org/10.1002/aur.2357

Marcinkiewcz, C. A., Mazzone, C. M., D’Agostino, G., Halladay, L. R., Hardaway, J. A., DiBerto, J. F., Navarro, M., Burnham, N., Cristiano, C., Dorrier, C. E., Tipton, G. J., Ramakrishnan, C., Kozicz, T., Deisseroth, K., Thiele, T. E., McElligott, Z. A., Holmes, A., Heisler, L. K., & Kash, T. L. (2016). Serotonin engages an anxiety and fear-promoting circuit in the extended amygdala. Nature, 537(7618), 97–101. https://doi.org/10.1038/nature19318

Marshall, C. R., Noor, A., Vincent, J. B., Lionel, A. C., Feuk, L., Skaug, J., Shago, M., Moessner, R., Pinto, D., Ren, Y., Thiruvahindrapduram, B., Fiebig, A., Schreiber, S., Friedman, J., Ketelaars, C. E. J., Vos, Y. J., Ficicioglu, C., Kirkpatrick, S., Nicolson, R.,… Scherer, S. W. (2008). Structural Variation of Chromosomes in Autism Spectrum Disorder. The American Journal of Human Genetics, 82(2), 477–488. https://doi.org/10.1016/j.ajhg.2007.12.009

Matson, J. L., & Cervantes, P. E. (2014). Commonly studied comorbid psychopathologies among persons with autism spectrum disorder. Research in Developmental Disabilities, 35(5), 952–962. https://doi.org/10.1016/j.ridd.2014.02.012

Mobbs, D., Yu, R., Rowe, J. B., Eich, H., FeldmanHall, O., & Dalgleish, T. (2010). Neural activity associated with monitoring the oscillating threat value of a tarantula. Proceedings of the National Academy of Sciences, 107(47), 20582–20586. https://doi.org/10.1073/pnas.1009076107

Olff, M. (2017). Sex and gender differences in post-traumatic stress disorder: an update. European Journal of Psychotraumatology, 8(sup4), 1351204. https://doi.org/10.1080/20008198.2017.1351204

Panzini, C. M., Ehlinger, D. G., Alchahin, A. M., Guo, Y., & Commons, K. G. (2017). 16p11.2 deletion syndrome mice perseverate with active coping response to acute stress – rescue by blocking 5-HT2A receptors. Journal of Neurochemistry, 143(6), 708–721. https://doi.org/10.1111/jnc.14227

Penzo, M. A., Robert, V., & Li, B. (2014). Fear Conditioning Potentiates Synaptic Transmission onto Long-Range Projection Neurons in the Lateral Subdivision of Central Amygdala. The Journal of Neuroscience, 34(7), 2432–2437. https://doi.org/10.1523/jneurosci.4166-13.2014

Penzo, M. A., Robert, V., Tucciarone, J., Bundel, D. D., Wang, M., Aelst, L. V., Darvas, M., Parada, L. F., Palmiter, R. D., He, M., Huang, Z. J., & Li, B. (2015). The paraventricular thalamus controls a central amygdala fear circuit. Nature, 519(7544), 455. https://doi.org/10.1038/nature13978

Portmann, T., Yang, M., Mao, R., Panagiotakos, G., Ellegood, J., Dolen, G., Bader, P. L., Grueter, B. A., Goold, C., Fisher, E., Clifford, K., Rengarajan, P., Kalikhman, D., Loureiro, D., Saw, N. L., Zhengqui, Z., Miller, M. A., Lerch, J. P., Henkelman, R. M.,… Dolmetsch, R. E. (2014). Behavioral Abnormalities and Circuit Defects in the Basal Ganglia of a Mouse Model of 16p11.2 Deletion Syndrome. Cell Reports, 7(4), 1077–1092. https://doi.org/10.1016/j.celrep.2014.03.036

Postorino, V., Kerns, C. M., Vivanti, G., Bradshaw, J., Siracusano, M., & Mazzone, L. (2017). Anxiety Disorders and Obsessive-Compulsive Disorder in Individuals with Autism Spectrum Disorder. Current Psychiatry Reports, 19(12), 92. https://doi.org/10.1007/s11920-017-0846-y

Pucilowska, J., Vithayathil, J., Tavares, E. J., Kelly, C., Karlo, J. C., & Landreth, G. E. (2015). The 16p11.2 Deletion Mouse Model of Autism Exhibits Altered Cortical Progenitor Proliferation and Brain Cytoarchitecture Linked to the ERK MAPK Pathway. The Journal of Neuroscience, 35(7), 3190–3200. https://doi.org/10.1523/jneurosci.4864-13.2015

Rein, B., & Yan, Z. (2020). 16p11.2 Copy Number Variations and Neurodevelopmental Disorders. Trends in Neurosciences. https://doi.org/10.1016/j.tins.2020.09.001

Roesch, M. R., Esber, G. R., Li, J., Daw, N. D., & Schoenbaum, G. (2012). Surprise! Neural correlates of Pearce–Hall and Rescorla–Wagner coexist within the brain. European Journal of Neuroscience, 55(7), 1190–1200. https://doi.org/10.1111/j.1460-9568.2011.07986.x

Rynkiewicz, A., & Łucka, I. (2018). Autism spectrum disorder (ASD) in girls. Co-occurring psychopathology. Sex differences in clinical manifestation. Psychiatria Polska, 52(4), 629–639. https://doi.org/10.12740/pp/onlinefirst/58837

Rynkiewicz, A., Schuller, B., Marchi, E., Piana, S., Camurri, A., Lassalle, A., & Baron-Cohen, S. (2016). An investigation of the ‘female camouflage effect’ in autism using a computerized ADOS-2 and a test of sex/gender differences. Molecular Autism, 7(1), 10. https://doi.org/10.1186/s13229-016-0073-0

Sanders, S. J., Ercan-Sencicek, A. G., Hus, V., Luo, R., Murtha, M. T., Moreno-De-Luca, D., Chu, S. H., Moreau, M. P., Gupta, A. R., Thomson, S. A., Mason, C. E., Bilguvar, K., Celestino-Soper, P. B. S., Choi, M., Crawford, E. L., Davis, L., Davis Wright, N. R., Dhodapkar, R. M., DiCola, M.,… State, M. W. (2011). Multiple Recurrent De Novo CNVs, Including Duplications of the 7q11.23 Williams Syndrome Region, Are Strongly Associated with Autism. Neuron, 70(5), 863–885. https://doi.org/10.1016/j.neuron.2011.05.002

Schumann, C. M., Hamstra, J., Goodlin-Jones, B. L., Lotspeich, L. J., Kwon, H., Buonocore, M. H., Lammers, C. R., Reiss, A. L., & Amaral, D. G. (2004). The Amygdala Is Enlarged in Children But Not Adolescents with Autism; the Hippocampus Is Enlarged at All Ages. The Journal of Neuroscience, 24(28), 6392–6401. https://doi.org/10.1523/jneurosci.1297-04.2004

Schwartz, C. E., & Neri, G. (2012). Autism and intellectual disability: Two sides of the same coin. American Journal of Medical Genetics Part C: Seminars in Medical Genetics, 160C(2), 89–90. https://doi.org/10.1002/ajmg.c.31329

Sebat, J., Lakshmi, B., Malhotra, D., Troge, J., Lese-Martin, C., Walsh, T., Yamrom, B., Yoon, S., Krasnitz, A., Kendall, J., Leotta, A., Pai, D., Zhang, R., Lee, Y.-H., Hicks, J., Spence, S. J., Lee, A. T., Puura, K., Lehtimäki, T.,… Wigler, M. (2007). Strong Association of De Novo Copy Number Mutations with Autism. Science, 316(5823), 445–449. https://doi.org/10.1126/science.1138659

Shackman, A. J., & Fox, A. S. (2016). Contributions of the Central Extended Amygdala to Fear and Anxiety. The Journal of Neuroscience, 36(31), 8050–8063. https://doi.org/10.1523/jneurosci.0982-16.2016

Shinawi, M., Liu, P., Kang, S.-H. L., Shen, J., Belmont, J. W., Scott, D. A., Probst, F. J., Craigen, W. J., Graham, B. H., Pursley, A., Clark, G., Lee, J., Proud, M., Stocco, A., Rodriguez, D. L., Kozel, B. A., Sparagana, S., Roeder, E. R., McGrew, S. G.,… Lupski, J. R. (2010). Recurrent reciprocal 16p11.2 rearrangements associated with global developmental delay, behavioural problems, dysmorphism, epilepsy, and abnormal head size. Journal of Medical Genetics, 47(5), 332. https://doi.org/10.1136/jmg.2009.073015

Solomon, M., Miller, M., Taylor, S. L., Hinshaw, S. P., & Carter, C. S. (2012). Autism Symptoms and Internalizing Psychopathology in Girls and Boys with Autism Spectrum Disorders. Journal of Autism and Developmental Disorders, 42(1), 48–59. https://doi.org/10.1007/s10803-011-1215-z

Sparks, B. F., Friedman, S. D., Shaw, D. W., Aylward, E. H., Echelard, D., Artru, A. A., Maravilla, K. R., Giedd, J. N., Munson, J., Dawson, G., & Dager, S. R. (2002). Brain structural abnormalities in young children with autism spectrum disorder. Neurology, 59(2), 184–192. https://doi.org/10.1212/wnl.59.2.184

Steinman, K. J., Spence, S. J., Ramocki, M. B., Proud, M. B., Kessler, S. K., Marco, E. J., Snyder, L. G., D’Angelo, D., Chen, Q., Chung, W. K., Sherr, E. H., & Consortium, S. V. (2016). 16p11.2 deletion and duplication: Characterizing neurologic phenotypes in a large clinically ascertained cohort. American Journal of Medical Genetics Part A, 170(11), 2943–2955. https://doi.org/10.1002/ajmg.a.37820

Szatmari, P., Liu, X., Goldberg, J., Zwaigenbaum, L., Paterson, A. D., Woodbury-Smith, M., Georgiades, S., Duku, E., & Thompson, A. (2012). Sex differences in repetitive stereotyped behaviors in autism: Implications for genetic liability. American Journal of Medical Genetics Part B: Neuropsychiatric Genetics, 159B(1), 5–12. https://doi.org/10.1002/ajmg.b.31238

Tian, D., Stoppel, L. J., Heynen, A. J., Lindemann, L., Jaeschke, G., Mills, A. A., & Bear, M. F. (2015). Contribution of mGluR5 to pathophysiology in a mouse model of human chromosome 16p11.2 microdeletion. Nature Neuroscience, 18(2), 182–184. https://doi.org/10.1038/nn.3911

Tolin, D. F., & Foa, E. B. (2006). Sex Differences in Trauma and Posttraumatic Stress Disorder: A Quantitative Review of 25 Years of Research. Psychological Bulletin, 132(6), 959–992. https://doi.org/10.1037/0033-2909.132.6.959

Tonnsen, B. L., Boan, A. D., Bradley, C. C., Charles, J., Cohen, A., & Carpenter, L. A. (2016). Prevalence of Autism Spectrum Disorders Among Children With Intellectual Disability. American Journal on Intellectual and Developmental Disabilities, 121(6), 487–500. https://doi.org/10.1352/1944-7558-121.6.487

Tovote, P., Fadok, J. P., & Lüthi, A. (2015). Neuronal circuits for fear and anxiety. Nature Reviews Neuroscience, 16(6), 317–331. https://doi.org/10.1038/nrn3945

Varodayan, F. P., Khom, S., Patel, R. R., Steinman, M. Q., Hedges, D. M., Oleata, C. S., Homanics, G. E., Roberto, M., & Bajo, M. (2018). Role of TLR4 in the Modulation of Central Amygdala GABA Transmission by CRF Following Restraint Stress. Alcohol and Alcoholism, 53(6), 642–649. https://doi.org/10.1093/alcalc/agx114

Varodayan, F. P., Minnig, M. A., Steinman, M. S., Oleata, C. S., Riley, M. W., Sabino, V., & Roberto, M. (2019). PACAP regulation of central amygdala GABAergic synapses is altered by restraint stress. Neuropharmacology, 107752. https://doi.org/10.1016/j.neuropharm.2019.107752

Wager, D., T., Barrett, F., L., Bliss-Moreau, E., Lindquist, A., K., Duncan, S., Kober, H., Joseph, J., Davidson, M., Mize, &, J. (2008). The neuroimaging of emotion. In M. Lewis, J. M. Haviland-Jones, & L. F. Barrett (Eds.), Handbook of emotions. The Guilford Press.

Walker, D. L., & Davis, M. (2008). Role of the extended amygdala in short-duration versus sustained fear: a tribute to Dr. Lennart Heimer. Brain Structure and Function, 213(1-2), 29–42. https://doi.org/10.1007/s00429-008-0183-3

Walsh, J. J., Christoffel, D. J., Heifets, B. D., Ben-Dor, G. A., Selimbeyoglu, A., Hung, L. W., Deisseroth, K., & Malenka, R. C. (2018). 5-HT release in nucleus accumbens rescues social deficits in mouse autism model. Nature, 560(7720), 589–594. https://doi.org/10.1038/s41586-018-0416-4

Weiss, L. A., Shen, Y., Korn, J. M., Arking, D. E., Miller, D. T., Fossdal, R., Saemundsen, E., Stefansson, H., Ferreira, M. A. R., Green, T., Platt, O. S., Ruderfer, D. M., Walsh, C. A., Altshuler, D., Chakravarti, A., Tanzi, R. E., Stefansson, K., Santangelo, S. L., Gusella, J. F.,… Consortium, A. (2008). Association between Microdeletion and Microduplication at 16p11.2 and Autism. The New England Journal of Medicine, 358(7), 667–675. https://doi.org/10.1056/nejmoa075974

Werling, D. M., & Geschwind, D. H. (2013). Sex differences in autism spectrum disorders. Current Opinion in Neurology, 26(2), 146–153. https://doi.org/10.1097/wco.0b013e32835ee548

White, S. W., Oswald, D., Ollendick, T., & Scahill, L. (2009). Anxiety in children and adolescents with autism spectrum disorders. Clinical Psychology Review, 29(3), 216–229. https://doi.org/10.1016/j.cpr.2009.01.003

Yang, M., Lewis, F. C., Sarvi, M. S., Foley, G. M., & Crawley, J. N. (2015). 16p11.2 Deletion mice display cognitive deficits in touchscreen learning and novelty recognition tasks. Learning & Memory, 22(12), 622–632. https://doi.org/10.1101/lm.039602.115

Yang, M., Mahrt, E. J., Lewis, F., Foley, G., Portmann, T., Dolmetsch, R. E., Portfors, C. V., & Crawley, J. N. (2015). 16p11.2 Deletion Syndrome Mice Display Sensory and Ultrasonic Vocalization Deficits During Social Interactions. Autism Research, 8(5), 507–521. https://doi.org/10.1002/aur.1465

Yu, K., Ahrens, S., Zhang, X., Schiff, H., Ramakrishnan, C., Fenno, L., Deisseroth, K., Zhao, F., Luo, M.-H., Gong, L., He, M., Zhou, P., Paninski, L., & Li, B. (2017). The central amygdala controls learning in the lateral amygdala. Nature Neuroscience, 20(12), 1680–1685. https://doi.org/10.1038/s41593-017-0009-9

Yu, K., Silva, P. G. da, Albeanu, D. F., & Li, B. (2016). Central Amygdala Somatostatin Neurons Gate Passive and Active Defensive Behaviors. The Journal of Neuroscience, 36(24), 6488–6496. https://doi.org/10.1523/jneurosci.4419-15.2016

Zimprich, A., Garrett, L., Deussing, J. M., Wotjak, C. T., Fuchs, H., Gailus-Durner, V., Angelis, M. H. de, Wurst, W., & Hölter, S. M. (2014). A robust and reliable non-invasive test for stress responsivity in mice. Frontiers in Behavioral Neuroscience, 8, 125. https://doi.org/10.3389/fnbeh.2014.00125

Zufferey, F., Sherr, E. H., Beckmann, N. D., Hanson, E., Maillard, A. M., Hippolyte, L., Macé, A., Ferrari, C., Kutalik, Z., Andrieux, J., Aylward, E., Barker, M., Bernier, R., Bouquillon, S., Conus, P., Delobel, B., Faucett, W. A., Goin-Kochel, R. P., Grant, E.,… Consortium, 16p11 2 European. (2012). A 600 kb deletion syndrome at 16p11.2 leads to energy imbalance and neuropsychiatric disorders. Journal of Medical Genetics, 49(10), 660. https://doi.org/10.1136/jmedgenet-2012-101203

